# Assessment of Antibiotic Resistance Profiles in Cultivable Coliform Organisms Isolated from Ganga River Waters Across the Upper, Middle, and Lower Ganga Stretch

**DOI:** 10.1101/2024.03.20.585858

**Authors:** Acharya Balkrishna, Shelly Singh, Sourav Ghosh, Ved Priya Arya, Mohini

## Abstract

Antibiotic resistance in coliform organisms isolated from the Ganga River waters across its Upper, Middle, and Lower stretches was assessed. The study revealed varying levels of antibiotic resistance among the coliform isolates, with differences observed across the different stretches of the river. In the Upper stretch, sites like Devprayag and Gangotri exhibited high levels of E. coli contamination, with corresponding moderate antibiotic resistance (MAR) values. In the Middle stretch, sites like Farrukhabad and Bithoor showed lower E. coli contamination compared to the Upper stretch but had higher MAR values. The Lower stretch, particularly sites like Revelganj and Bahachoki in Bihar, displayed significantly elevated E. coli contamination levels and MAR values. These findings underscore the presence of antibiotic-resistant coliform organisms in different segments of the Ganga River, emphasizing the need for targeted interventions to mitigate antibiotic resistance spread in this vital water resource. Understanding the antibiotic resistance profiles at specific sites along the Ganga River can inform strategies to preserve water quality and protect public health in regions dependent on this iconic river.

## Introduction

With its profound cultural, religious, and economic significance in India, the Ganga River stands as a crucial water source for domestic, industrial, and agricultural purposes. Given its pivotal role, understanding and addressing threats to the river’s water quality, such as antibiotic resistance, become imperative (Ali et al., 2021). The escalating global challenge of antibiotic resistance, fuelled by indiscriminate usage across various sectors, poses a significant public health threat. The Ganga River, heavily impacted by human activities, emerges as a potential hotspot for disseminating antibiotic resistance. Hence, assessing the antibiotic resistance profiles of coliform organisms in this iconic river is critical for comprehending and mitigating this growing concern (Rout et al., 2023). Despite the Ganga River’s paramount importance, a notable gap exists in comprehensive studies addressing the antibiotic resistance profiles of coliform organisms within its waters. This research endeavor seeks to fill this void by presenting an in-depth analysis of the upper, middle, and lower stretches of the ganga river, thereby enriching the existing knowledge base (Singh et al., 2020). The consequences of antimicrobial-resistant infections extend beyond their challenging and sometimes insurmountable treatment. Such infections result in prolonged illnesses, heightened healthcare costs, and elevated mortality rates, transforming once easily treatable infections into life-threatening conditions. The global surge in antibiotic resistance constitutes a formidable threat to public health, diminishing the efficacy of common antibiotics against widespread bacterial infections. This impact is particularly felt in low-resource settings and vulnerable populations, where the effective treatment of common infections becomes increasingly challenging.

Furthermore, antibiotic residues and resistant bacteria in the environment are recognized as environmental pollutants that can influence bacterial populations, foster resistance spread, and induce adverse ecological effects. This environmental contamination can, in turn, affect human health through multiple pathways, including contaminated water sources and exposure through the food chain. The antibiotic-resistant bacteria present in the environment may harbor pathogenic potential for humans, posing a direct risk of causing infectious diseases that are challenging to treat (Svetlana Iuliana Polianciuc et al., 2020). The issue of antibiotic pollution is multifaceted, originating from both the excretion of antibiotics by humans and animals, as well as the improper disposal of unused or expired antibiotics. Antibiotics administered to humans and animals are excreted unchanged in feces and urine, contributing to the introduction of these pharmaceuticals into the environment through sewage systems and wastewater treatment plants (Larsson & Flach, 2021). Simultaneously, the improper disposal of antibiotics further exacerbates environmental contamination, manifesting in direct contamination in aquaculture, plant production, and the release of antibiotics via waste streams from the production of antibiotics. Animal farms, renowned for generating substantial quantities of waste, emerge as significant contributors to antibiotic pollution. This waste, laden with antibiotics and antibiotic-resistant bacteria, poses a substantial risk as it permeates soil, water, and air, thereby facilitating the dissemination of antibiotic resistance genes. The magnitude of the problem is further amplified by the discharge of wastewater from antibiotic manufacturing sites, which often contain elevated concentrations of antibiotics. Illustratively, a wastewater treatment plant in India, servicing waste from 90 drug manufacturers, was found to release 45 kg of ciprofloxacin into a nearby river daily. The healthcare sector is not exempt from contributing to antibiotic pollution, with hospitals and municipalities discharging wastewater containing both antibiotics and antibiotic-resistant bacteria. This discharge further propagates the contamination of soil, water, and air, intensifying the dissemination of antibiotic-resistance genes. Moreover, antibiotic pollution can be attributed to industrial activities such as the manufacturing of antibiotics or the inadequate disposal of waste generated during pharmaceutical production. In essence, the diverse sources of antibiotic pollution underscore the urgent need for comprehensive strategies to mitigate this environmental threat.

The gravity of the situation is underscored by the fact that antimicrobial resistance directly contributed to 1.27 million global deaths in 2019. The misuse and overuse of antimicrobials in humans, animals, and plants emerge as primary drivers in developing drug-resistant pathogens. Addressing these challenges in the context of the Ganga River’s ecosystem is not only a matter of environmental concern but also a critical facet of safeguarding public health and ensuring sustainable water resources for future generations.

## Methodology

### Study area

The Ganga River Basin, stretching over a remarkable distance of approximately 2,525 kilometers from Gaumukh to Ganga Sagar, is the focal point for our comprehensive research on antibiotic resistance . The catchment area of the Ganga River System is about 861,404 square kilometers, covering parts of Uttarakhand and Uttar Pradesh states (Santy et al., 2020; Singh et al., 2022). Approximately 43% of India’s population lives in the Ganga basin, which covers 26.3% of the country’s total geographical area. The river Ganga passes along 29 Class I cities, 23 Class II cities, and approximately 50 towns.

Within this vast expanse, the upper stretch from Gaumukh to Narora epitomizes the river’s purity, as it meanders through the untouched landscapes of Uttarakhand and Uttar Pradesh. This region serves as a crucial benchmark for understanding the baseline antibiotic resistance levels in natural environments. The middle stretch from Badaun to Ballia presents a transition zone, where the impacts of human activities begin to manifest. Sampling sites along this stretch, such as Bijnor and Narora, provide insights into the escalating anthropogenic pressures on water quality and antibiotic pollution. Moving downstream, the lower stretch from Revelganj to Gangasagar encompasses highly populated regions where urbanization and industrialization exert significant stress on the river ecosystem. Sites like Revelganj, Patna, and Gangasagar offer critical data points for assessing the cumulative effects of human influence on antibiotic resistance within the Ganga River Basin.

**Figure 1.**
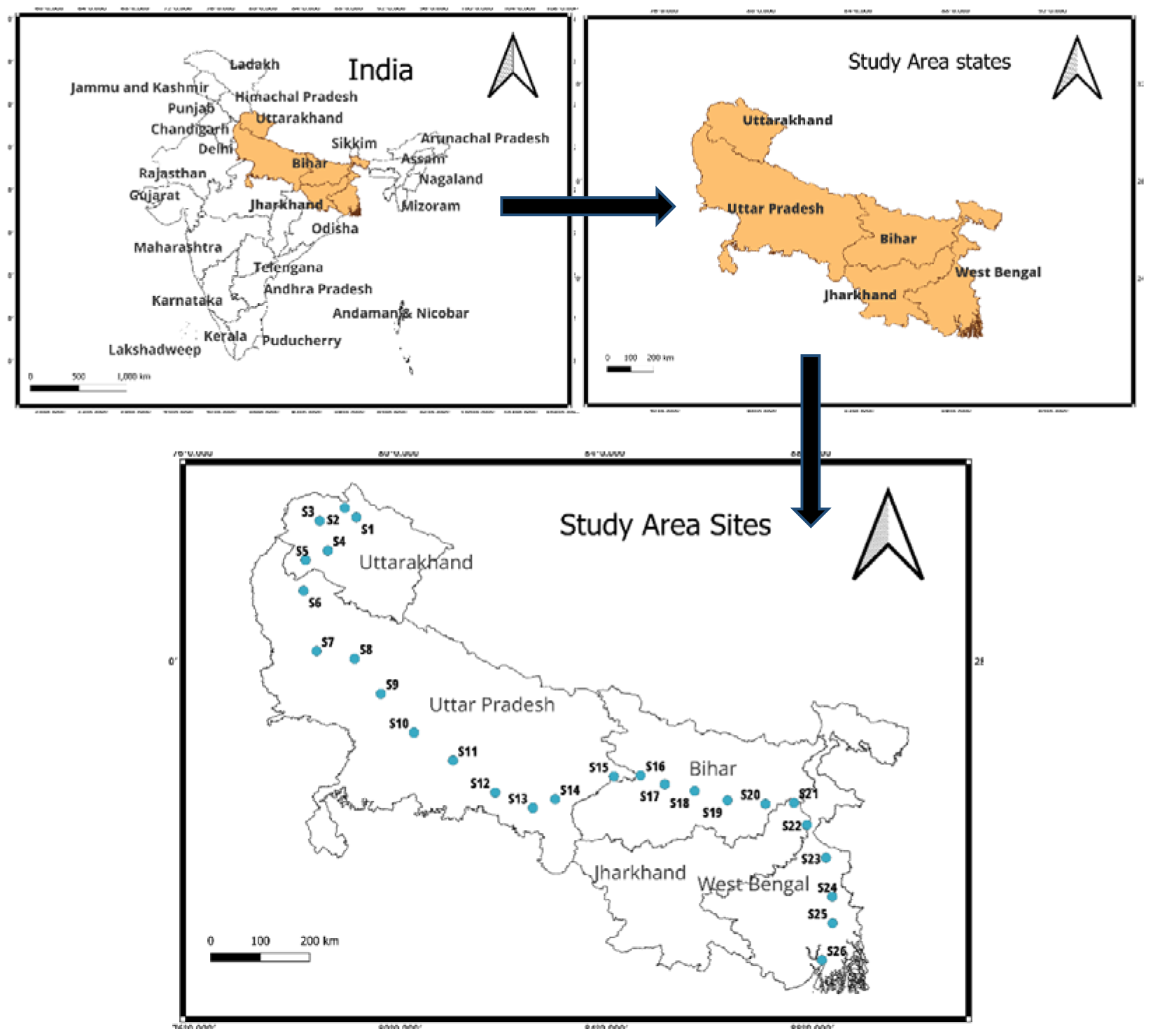
Location of the study area, i.e., 26 sites of UK, UP, Bihar, Jharkhand, and West Bengal, India.

### Sampling of Water samples

A systematic sampling approach ensures accurate representation and adherence to regulatory standards. A pre-sterilized 20 mL glass container is utilized during the collection process, with the utmost care to wear gloves to prevent contamination. Samples are drawn from a depth between 20 and 30 centimeters below the water’s surface, capturing a representative snapshot of microbial content. The containers are meticulously filled to the brim to provide sufficient volume for analysis. To maintain sample integrity, they are promptly stored in a refrigerated container filled with ice, ensuring a constant temperature range of 2-4 degrees Celsius during transportation. This meticulous approach not only aligns with the guidelines set by the Central Pollution Control Board (CPCB) but also adheres to the 2012 standards of the American Public Health Association (APHA).

### Cultivation of faecal coliform

In this study, the cultivation of faecal coliform using selective media and the spread plate method was employed as a fundamental technique for quantifying faecal coliform presence in water. Selective media Eosin Methylene Blue agar is chosen for its ability to inhibit the growth of non-coliform bacteria while facilitating the growth of faecal coliforms, particularly Escherichia coli. The spread plate method involved spreading a known volume (100uL) of the sample onto the surface of the selective agar plate using a sterile spreader, followed by incubation at 35° C temperature. Colonies of *Escherichia coli* appeared as black with green metallic sheen plus faecal coliform bacteria appeared as distinctive pink or purple colonies on these agar plates due to their ability to ferment lactose.

### Selection of relevant antibiotics

### Antibiotic Susceptibility Test

The Disk Diffusion Method (Kirby-Bauer Test), which entails inserting antibiotic-impregnated paper disks on a culture plate infected with bacteria, was used to detect antibiotic sensitivity and resistance patterns using nutrient broth (HiMedia, M002), antibiotic disc, and micro titre plates. Following incubation at 37 °C for 24h, each disk’s turbidity is measured; a positive result indicates that the bacteria are resistant to the antibiotic (Bauer et al. 1996).

MAR (Multiple antibiotic resistance) indices for each isolate were then computed by dividing the number of antibiotics to which the isolate was resistant by the total number of antibiotics tested. In the context of a single isolate, the MAR index may be expressed as a/b, where a denotes the number of antibiotics to which the isolate exhibited resistance and b the number of antibiotics to which the isolate was exposed. A MAR index score of more than 0.2 is thought to have come from high-risk sources of contamination, such as commercial poultry farms, dairy cows, pigs, and humans, where antibiotics are often used. A MAR index value of less than or equal to 0.2 indicates that the strain originated from an animal handled with antibiotics seldom (Vivekanandhan et al, 2002; Watkinson et al, 2007; Krumperman 1983).

## Result and Discussion

The research aimed to assess the levels of faecal coliform or *E. coli* and antibiotic resistance (MAR index) in water samples collected from various sites along the Ganga River spanning five states in India: Uttarakhand, Uttar Pradesh, Bihar, Jharkhand, and West Bengal. The data obtained from the analysis are presented below in Table 3.

**Table 1.**
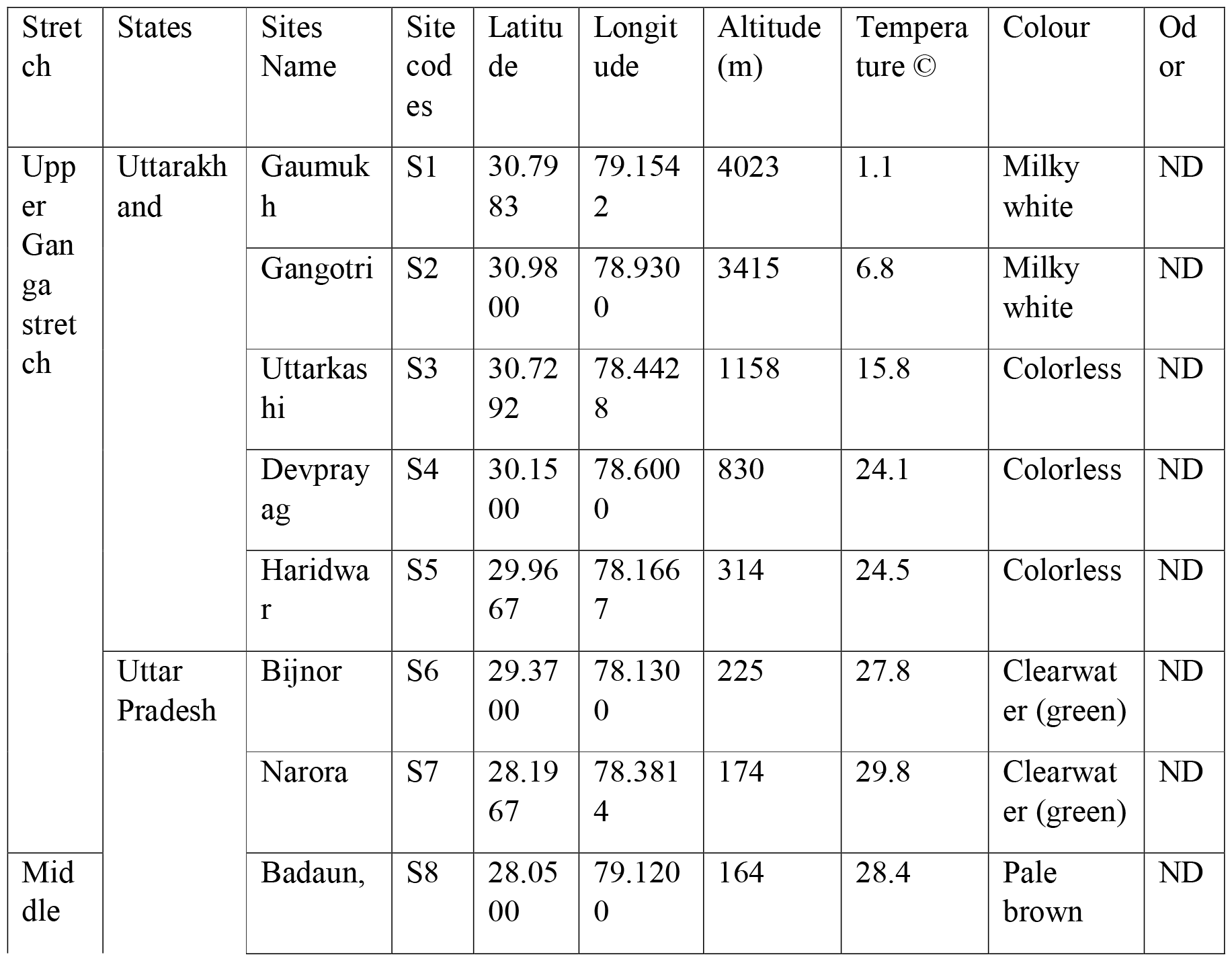
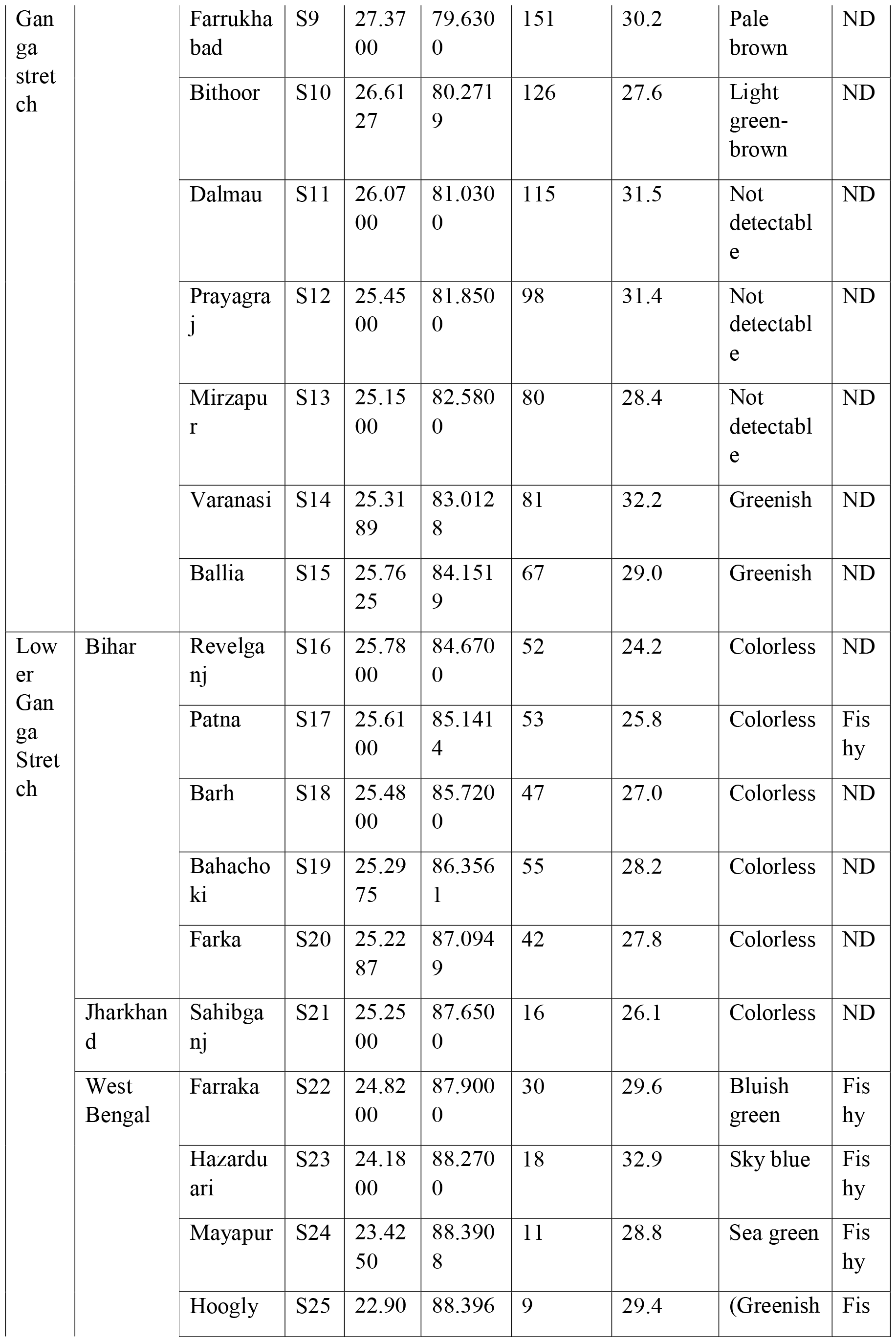
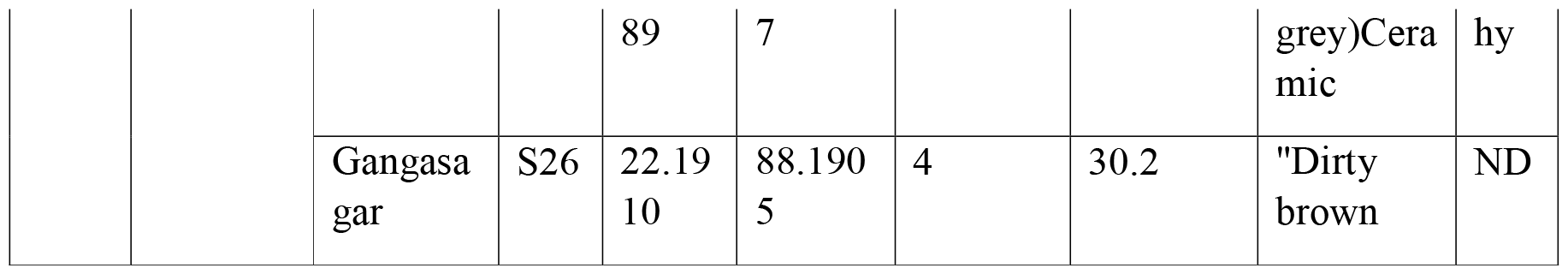
On-site recorded data of river Ganga of UK, UP, Bihar, Jharkhand, and West Bengal, India.

**Table 2.**
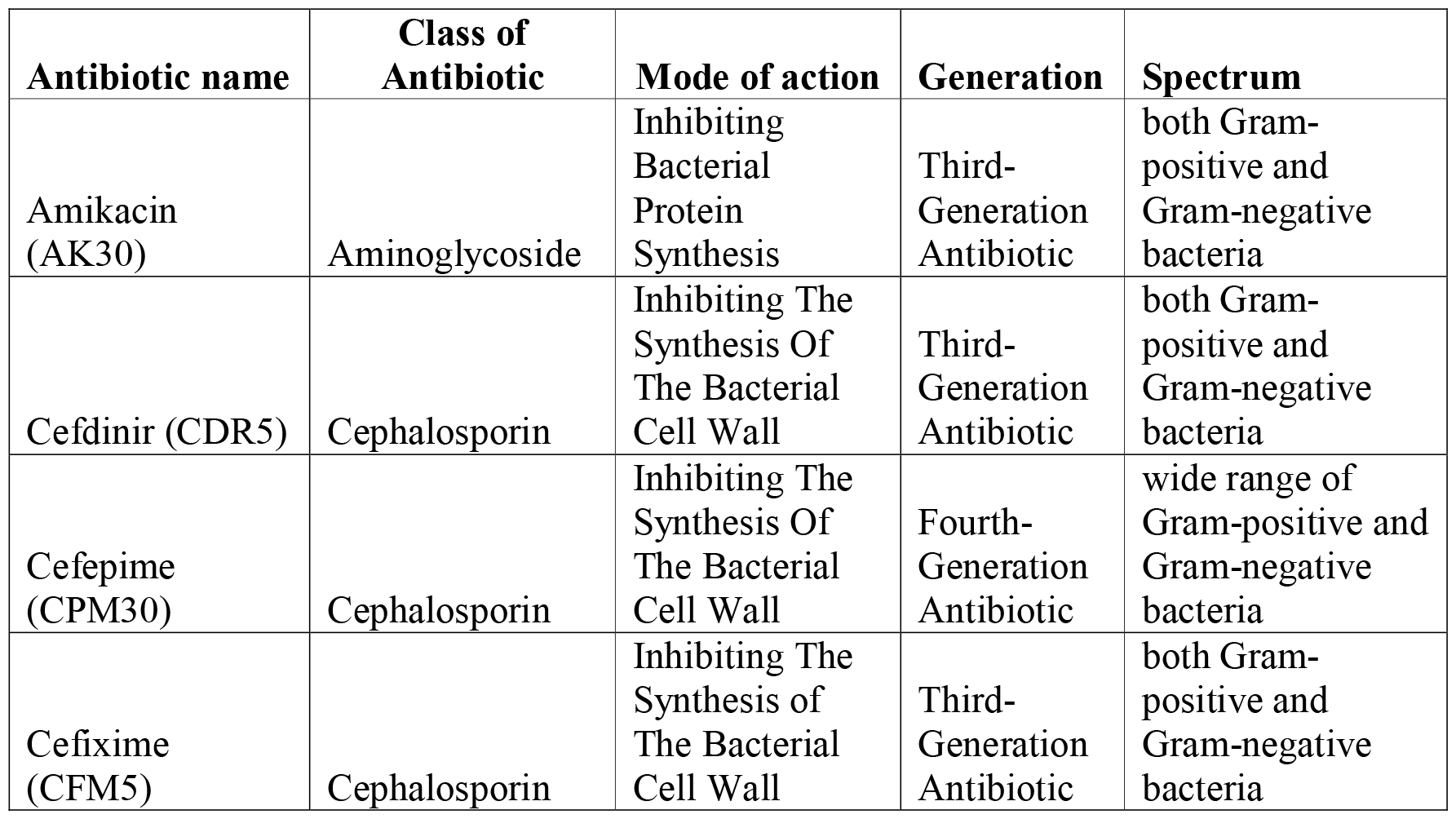
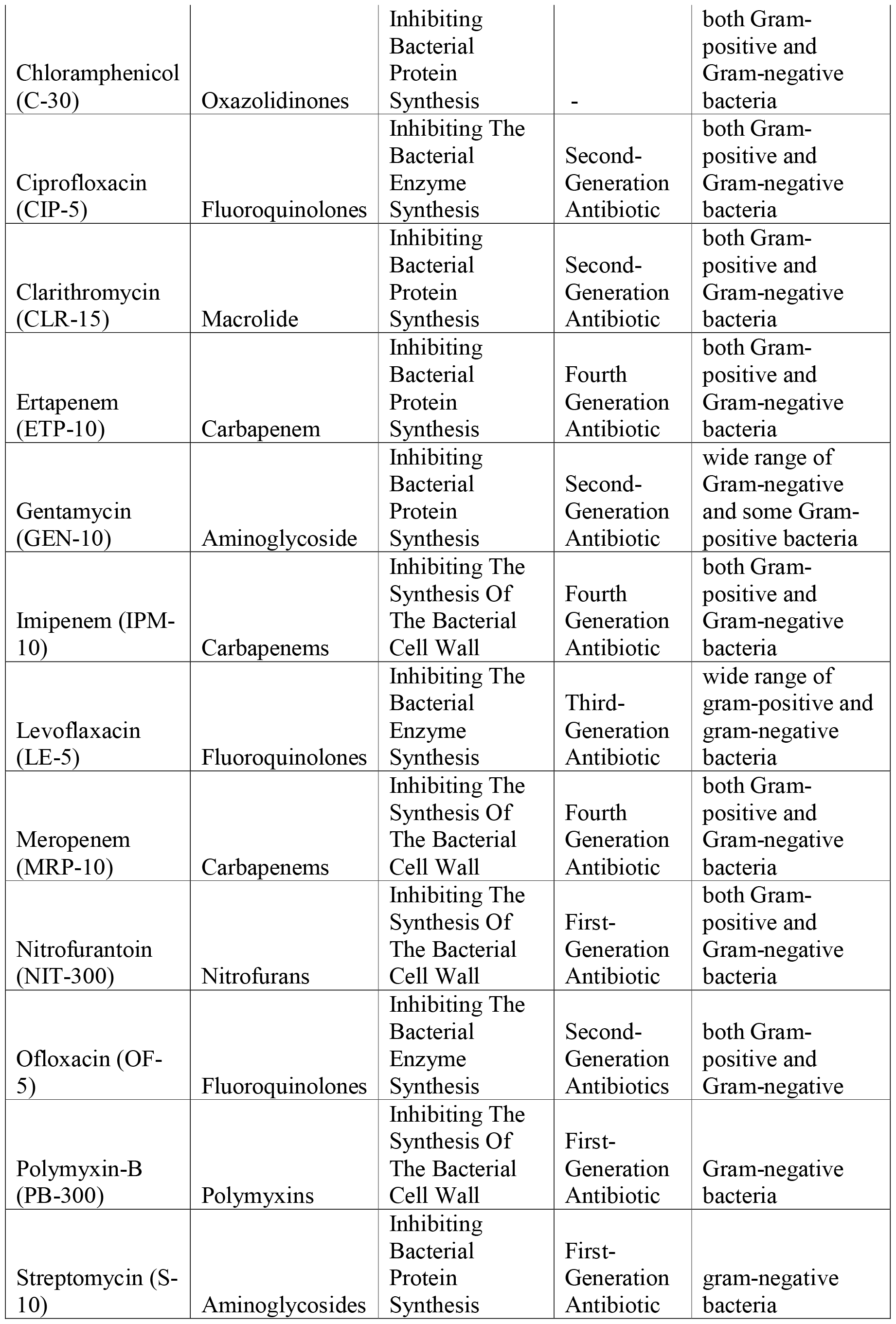
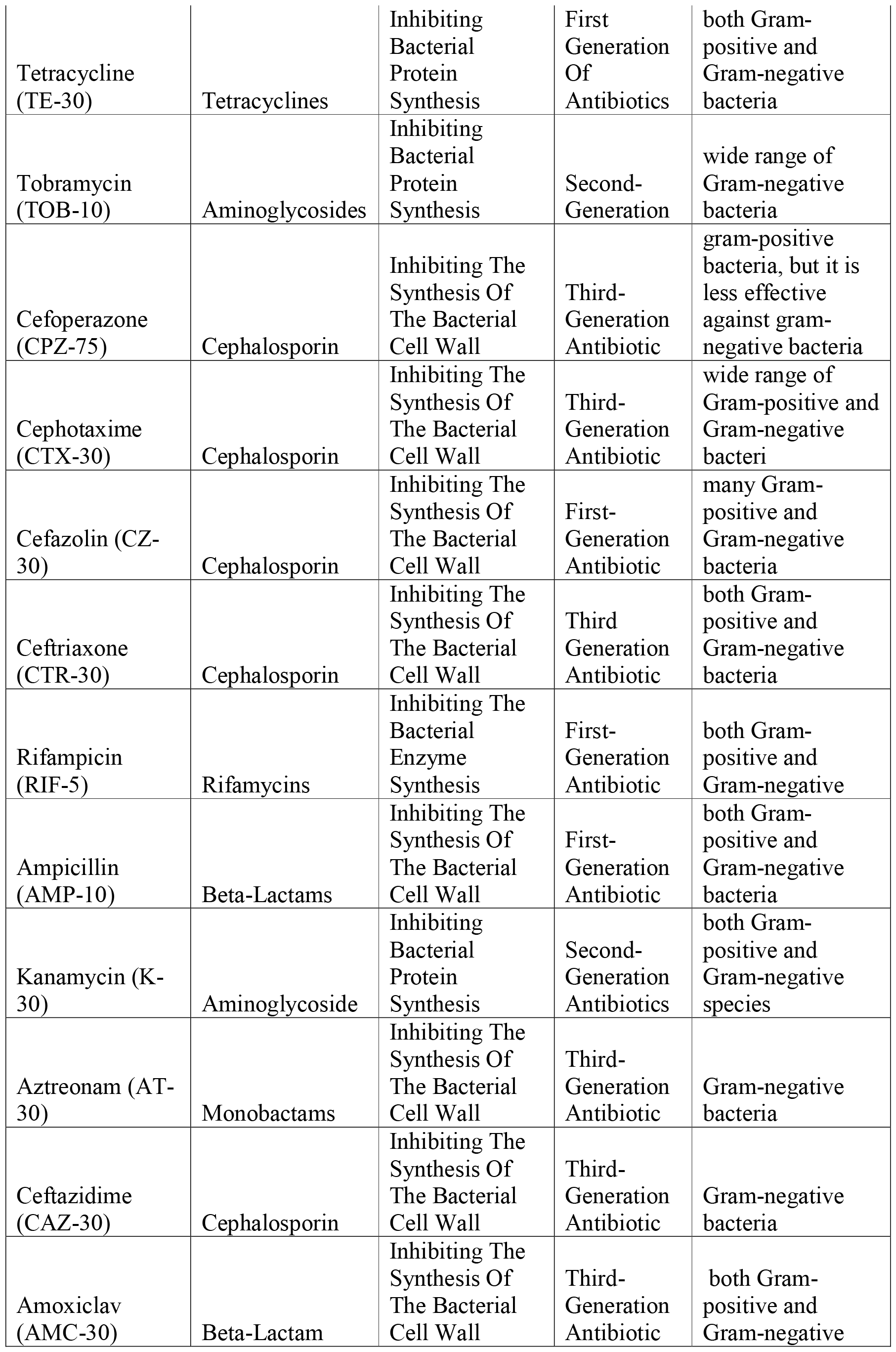
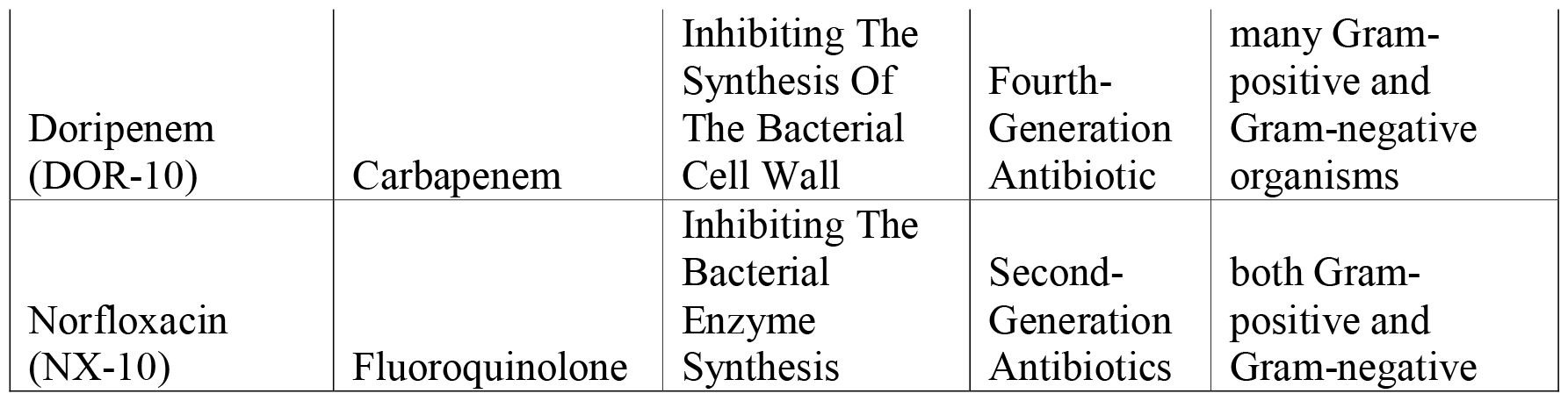
Below are the given selected 32 broad-spectrum antibiotic names, classes of antibiotics, mode of action, generation, and spectrum.

**Table 3.** Iindicates the presence of Total coliform and MAR values of river Ganga of UK, UP, Bihar, Jharkhand, and West Bengal, India.

| Stretch | States | Sites Name | Site codes | E coli Log 10 (Cfu/ml) | MAR |
| --- | --- | --- | --- | --- | --- |
| Upper stretch | Uttarakhand | Gaumukh | S1 | 0.00 | 0 |
|  |  | Gangotri, | S2 | 8.97 | 0.162 |
|  |  | Uttarkashi | S3 | 8.41 | 0 |
|  |  | Devprayag | S4 | 9.19 | 0.297 |
|  |  | Haridwar | S5 | 8.37 | 0.054 |
|  | Uttar Pradesh | Bijnor | S6 | 8.92 | 0.189 |
|  |  | Narora | S7 | 0.92 | 0.324 |
|  |  |  | Median | 8.41 | 0.16 |
| Middle stretch |  | Badaun, | S8 | 6.80 | 0.216 |
|  |  | Farrukhabad | S9 | 8.72 | 0.27 |
|  |  | Bithoor | S10 | 8.89 | 0.189 |
|  |  | Dalmau | S11 | 7.89 | 0.405 |
|  |  | Prayagraj | S12 | 7.92 | 0.459 |
|  |  | Mirzapur | S13 | 7.73 | 0.27 |
|  |  | Varanasi | S14 | 7.71 | 0.324 |
|  |  | Ballia | S15 | 8.30 | 0.243 |
|  |  |  | Median | 7.90 | 0.27 |
| Lower stretch | Bihar | Revelganj | S16 | 10.15 | 0.405 |
|  |  | Patna | S17 | 10.26 | 0.2162 |
|  |  | Barh | S18 | 9.21 | 0.27 |
|  |  | Bahachoki | S19 | 10.71 | 0.243 |
|  |  | Farka | S20 | 8.41 | 0.216 |
|  | Jharkhand | Sahibganj | S21 | 10.44 | 0.405 |
|  | West Bengal | Farraka | S22 | 3.35 | 0.2162 |
|  |  | Hazarduari | S23 | 2.92 | 0.378 |
|  |  | Mayapur | S24 | 9.06 | 0.378 |
|  |  | Hoogly | S25 | 9.36 | 0.378 |
|  |  | Gangasagar | S26 | 8.27 | 0.432 |
|  |  |  | Median | 9.21 | 0.38 |

Notably, the faecal coliform levels varied significantly across different sites and states, reflecting diverse sources of contamination along the river basin. In the upper stretches of the river basin, encompassing Uttarakhand and select sites in Uttar Pradesh, we observe varying levels of E. coli and Multi-Antibiotic Resistance (MAR), shedding light on microbial contamination dynamics. Notably, Gaumukh in Uttarakhand displays an absence of E. coli, indicating minimal contamination, alongside a MAR value of 0. Moving downstream to Gangotri, Uttarkashi, Devprayag, and Haridwar, E. coli concentrations gradually increase. Devprayag records the highest E. coli level among these sites, accompanied by a MAR value of 0.297. Additionally, sites such as Bijnor and Narora in Uttar Pradesh exhibit intermediate E. coli levels and MAR values. These findings underscore the complexity of microbial contamination and antibiotic resistance patterns in the upper stretches of the river basin, emphasizing the need for comprehensive monitoring and management strategies to safeguard water quality. Emphasizing the need for a multifaceted approach to river health management

In the middle stretches of the river basin, located primarily in Uttar Pradesh, we encounter a series of sites showcasing varied levels of E. coli and Multi-Antibiotic Resistance (MAR). These sites play a crucial role in understanding the microbial contamination dynamics within this region. Notably, sites such as Badaun, Farrukhabad, Bithoor, Dalmau, Prayagraj, Mirzapur, Varanasi, and Ballia exhibit diverse concentrations of E. coli. Among these, Dalmau records the highest level of E. coli. Moreover, MAR values differ across these sites, with Dalmau also displaying the highest MAR value. This comprehensive assessment highlights the complexity of microbial contamination and antibiotic resistance in the middle stretches of the river, underscoring the need for thorough environmental monitoring and management strategies to mitigate health risks.

In the lower stretches of the river basin, spanning Bihar, Jharkhand, and West Bengal, specific sites provide data on E. coli levels and Multi-Antibiotic Resistance (MAR), offering insights into microbial contamination dynamics. Bihar’s sites, including Revelganj, Patna, Barh, Bahachoki, and Farka, exhibit varying concentrations of E. coli, with Revelganj and Patna displaying notably high levels. Jharkhand’s Sahibganj site also presents elevated E. coli levels. In West Bengal, Farraka, Hazarduari, Mayapur, Hooghly, and Gangasagar show diverse E. coli concentrations, indicating variability across the region. These findings underscore the necessity for comprehensive environmental monitoring and management strategies to address microbial contamination and antibiotic resistance effectively within the lower stretches of the river basin.

Furthermore, the Multi-Antibiotic Resistance (MAR) index provided insights into the prevalence of antibiotic resistance among coliform bacteria in the water samples. The MAR index values ranged from 0 to 0.459, with higher values indicating increased resistance to multiple antibiotics.

**The antibiogram of total coliform/E. coli** isolated from the surface water of River Ganga across 26 sites spanning Uttarakhand, Uttar Pradesh, Bihar, Jharkhand, and West Bengal, India, revealed varying levels of antibiotic resistance and susceptibility. The percentage resistance and susceptibility of different antibiotics are summarized in the table 4 below:

**Table 4.** Antibiogram of total coliform\E.coli isolated from the surface water of river Ganga of UK, UP, Bihar, Jharkhand, and West Bengal, India.

| Antibiotic | Resistant | Sensitive |
| --- | --- | --- |
| (AK30) | 3.80% | 96.15% |
| (CDR5) | 84.60% | 15.4% |
| (CPM30) | 42.30% | 57.700% |
| (CFM5) | 65.30% | 34.70% |
| (C-30) | 3.80% | 96.15% |
| (CIP-5) | 3.80% | 96.15% |
| (CLR-15) | 42.30% | 57.700% |
| (ETP-10) | 23.07% | 76.90% |
| (GEN-10) | 3.80% | 96.15% |
| (IPM-10) | 15.38% | 84.62% |
| (LE-5) | 3.80% | 96.15% |
| (MRP-10) | 3.80% | 96.15% |
| (NIT-300) | 46.15% | 53.85% |
| (OF-5) | 0 | 100% |
| (PB-300) | 15.38% | 84.62% |
| (S-10) | 7.69% | 92.30% |
| (TE-30) | 0 | 100% |
| (TOB-10) | 7.69% | 92.30% |
| (CPZ-75) | 3.80% | 96.15% |
| (CTX-30) | 0 | 100% |
| (CZ-30) | 53.80% | 46.15% |
| (CTR-30) | 0 | 100% |
| (RIF-5) | 34.60% | 65.38% |
| (AMP-10) | 61.53% | 38.40% |
| (K-30) | 11.53% | 88.47% |
| (AT-30) | 38.46% | 61.53% |
| (CAZ-30) | 23.07% | 76.92% |
| (AMC-30) | 34.61% | 65.39% |
| (DOR-10) | 0 | 100% |
| (NX-10) | 0 | 100% |

## Conclusion

The information presented provides a detailed summary of antibiotic susceptibility and resistance percentages for various drugs. Notably, some antibiotics have high susceptibility rates, demonstrating their efficiency in eradicating bacterial infections without developing resistance. For example, OF-5, TE-30, CTX-30, CTR-30, DOR-10, and NX-10 are completely susceptible, with a 100% susceptibility rate and no resistance found. In contrast, other antibiotics confront significant resistance concerns, with high rates of bacterial resistance reported. Antibiotics such as CDR5, CFM5, NIT-300, RIF-5, and AMP-10 have high resistance rates, ranging from 34.60% to 84.60%. These findings highlight the vital relevance of cautious antibiotic usage and continuing surveillance in combating growing resistance trends and maintaining the efficacy of antibiotic therapies.

## Conflicts of Interest

### The authors declare that there are no conflicts of interest

#### CRediT authorship contribution statement

Sourav Ghosh: Writing – review & editing, Writing – original draft, Supervision, Project administration, Conceptualization. Acharya Balkrishna, Vedpriya Arya: Resources, Conceptualization. Shelly Singh: Methodology, Writing – review & editing

#### Financial Assistance

This research was supported by the National Mission for Clean Ganga, Ministry of Jal Shakti, Government of India under the Namami Gange Mission-II (Sanction order. F. No. Ad-35013/4/2022-KPMG-NMCG).

## Acknowledgments

The authors would acknowledge reverted Swami Ramdev Ji for his guidance during the execution of the study. The authors are thankful to the National Mission for Clean Ganga, Ministry of Jal Shakti for the effective execution of the project under the Namami Gange Mission-II. Further, the authors also thank the Ministry of AYUSH under Grant-in-Aid for the Establishment of the Centre of Excellence of Renovation and Upgradation of Patanjali Ayurveda Hospital, Haridwar, India. The authors also thankful to Mr. Harshit Thakur for preparation of Infographics for the manuscript.

## Notes

### Competing Interest Statement

The authors have declared no competing interest.

